# Understanding the Lipidome at the Systems Level with lipidomeR

**DOI:** 10.1101/2020.03.16.994061

**Authors:** Tommi Suvitaival, Cristina Legido-Quigley

## Abstract

Lipidomics is one of the fastest-growing areas of molecular profiling in medicine. While increasing amounts of lipidomics data are generated, tools for analyzing and interpreting these data are not equally widely available. We present the lipidomeR -- a tool specifically designed for systematic interpretation of large lipidome-wide studies. The lipidomeR binds together statistical analysis and high-dimensional visualization, providing a reproducible pipeline for rapid interpretation of the lipidome via integrative publication-ready figures. The lipidomeR package is available through the Comprehensive R Archive (CRAN).

We demonstrate the lipidomeR with three studies from the Metabolomics Workbench repository, ranging from the human plasma reference material to breast tumor tissue and to the progression of non-alcoholic liver disease (NAFLD) in the liver. In these studies, lipidomeR reveals a diversity of lipidomic patterns, both, within and between the lipid classes as well as over the stages of progression of the diseases.

## 1 Introduction

The development of mass-spectrometry and other omics technologies have made lipids an increasingly accessible target to measure and study. In relative terms, lipidomics is currently the fastest-growing field among the omics technologies.

Lipids have very diverse chemical structures but a large number of lipid molecules have one to three hydrophobic fatty acid chains attached to a hydrophilic head group. These lipids, which share the fatty acid chain structure, are classified by the type of the head group. For instance, triacylglycerols (TGs), which colloquially are known as triglycerides, have three fatty acid chains attached to a glycerol head group.

While lipids of the same class share the same structure type, individual species differentiate by the length of the fatty acid chains and by the level of saturation. The length of fatty acid chains is quantified by the number of carbon atoms and the inverse level of saturation is quantified by the number of unsaturated double bonds in the fatty acid chains. Lipid species differing by these characteristics can be identified with mass spectrometry.

There is growing evidence that the length and saturation of a lipid is associated with function. The structural relationship of lipid species in one class gives an opportunity to not only group the lipids by the class but also organize them based on the two characteristic properties of length and saturation. In this paper, we present a tool for presenting lipids in an organized map, where lipids are grouped by the class and organized on two-dimensional maps based on the length and saturation.

Two-dimensional presentations of lipids within one class have already been manually created and used for reporting lipidomics data, successfully providing new insights on lipid activity in health and disease (see, e.g., Hyysalo, et al., 2014^1^). Here, we provide a tool for visualizing the lipidome in this systematic manner. Further, we tap the visualization tool directly to the output of a high-throughput statistics inference engine^2^, making the analysis, visualization and interpretation of complex lipidomics data a seamless, fast and reproducible process.

The growth of lipidomics and its adoption in translational medical research and has led to an increasing demand in the tools for processing, analyzing, interpreting and integrating the data. To this date, the tools for interpreting the lipidome have to a large extent been limited to the following approaches or their variants: (1) the identification of individual markers based on univariate or regression model-based testing, as outlined previously, combined with a contextual interpretation based on a literature search of the identified markers (see, e.g., Rauschert, et al., 2016^3^); (2) the analysis of lipid class-specific total amounts or average levels (see, e.g., Stegemann, et al., 2011^4^); (3) the analysis of average levels in groups or clusters of correlated lipids (see, e.g., Huopaniemi, et al., 2010^5^); (4) the lipid class-specific regression analysis between the saturation of lipids to and their concentration (see, e.g., Rhee, et al. 2011^6^), (5) the interpretation of the lipid levels in the context of broader pathways derived from literature or other omics technologies, such as transcriptomics or proteomics (see, e.g., Tonks, et al., 2016^7^).

In this manuscript, we present a toolbox that gives a means to interpreting lipidomics data in a way that can be viewed as a combination of the categories 1, 2, and 4 from the above listing. Further, the package includes additional capabilities that can be useful for the categories 3 and 5. The toolbox leverages on established R packages and provides a pipeline for the generation of rich data analyses that present the lipidome in relation to explanatory variables of the study and present the lipidome in the context of structural information on the lipid class, size and level of saturation. We argue that these are the three key descriptors for understanding underlying mechanisms in the lipidome. On the other hand, understanding the influence of competing factors and their relative contributions is essential for understanding the extent and importance of an association, which is made possible with side-by-side visualizations of independent variables from the model.

## 2 Results

### 2.1. The lipidomeR Package

The lipidomeR is written in R^8^ (version 3.6.2) and is available through the Comprehensive R Archive Network (CRAN), which is the major R package repository provided by the R-project. The package is available for installation with the command ‘install.packages(“lipidomeR”)’ for R versions 3.5.0 or newer. Key back-end dependencies of the lipidomeR are the limma package^2^ for statistical inference, the ggplot2 package^9^ for the visualization and the stringr package^10^ for parsing and enumerating the lipid names into mappable values.

The workflow of a lipidomeR analysis is outlined in Figure 1. Briefly, it consists of (1) loading the data into R, (2) defining the list of lipid names, (3) enumerating the lipid names into mappable values of fatty acid chain length and saturation, (4) computing the regression models to explain the lipid observations based on the experiment design, and (5) creating the lipidomeR heatmap that integrates the model statistics over the measured lipidome. The enumeration of the lipid names (Step 3) is done using the ‘map_lipid_names()’ function of the lipidomeR package, The regression models (Step 4) are fitted using the ‘compute_models_with_limma()’ function, and, finally, the results of the models are visualized (Step 5) using the ‘heatmap_lipidome_with_limma()’ function. This most typical route of analysis is shown with bold lines from top to bottom in the middle-part of Figure 1 (see Section 2.3 for an example).

**Figure 1:**
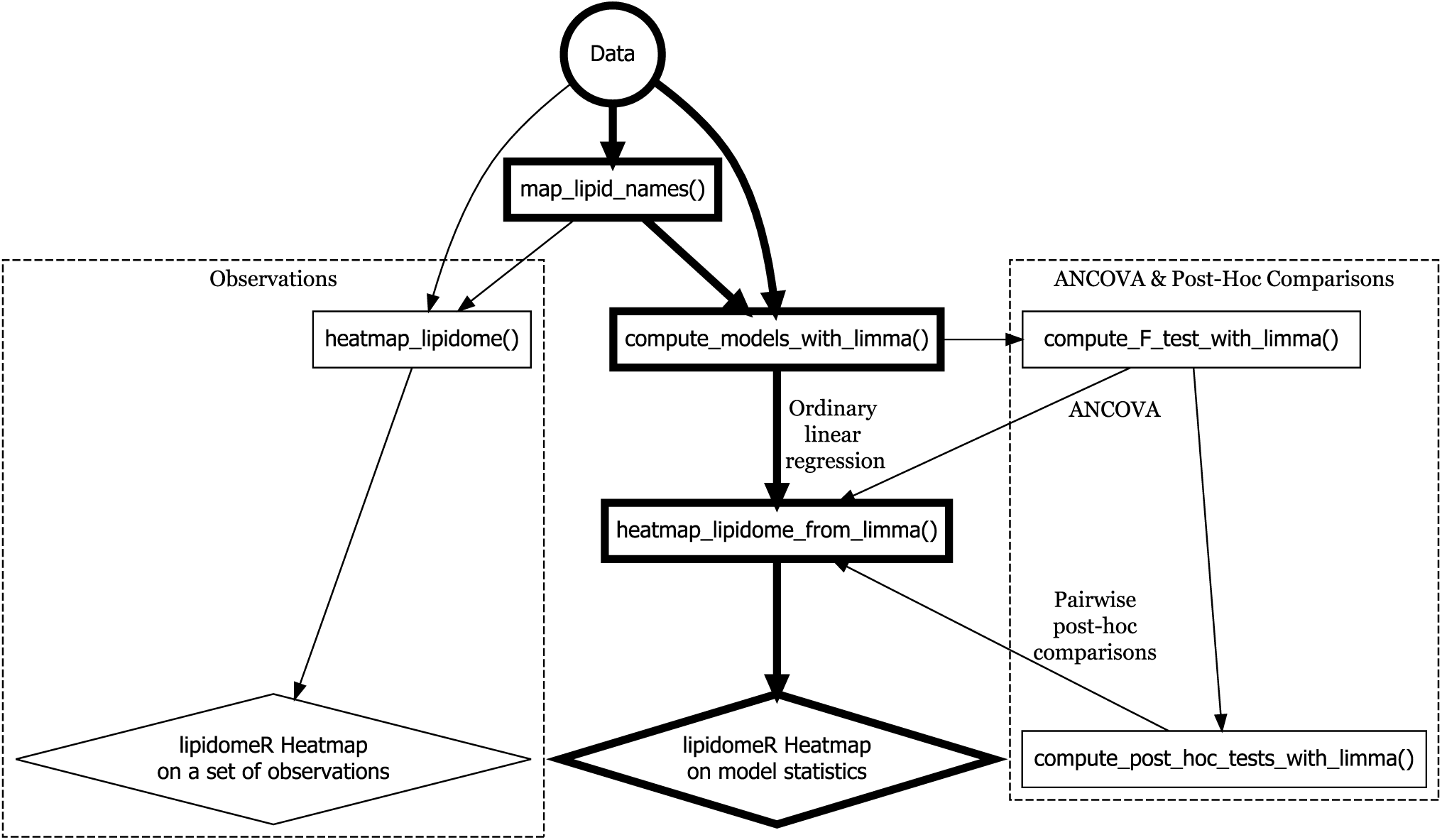
Workflow of a lipidomeR analysis from top (data) to bottom (result). The input (data) is shown as circle, the output (i.e., the result figures) are shown as diamonds and the functions of the lipidomeR package are shown as rectangles. The main route with an ordinary linear regression model is shown in bold in the middle. Alternative route with an analysis of covariance (ANCOVA) and pairwise post-hoc comparisons of significant lipids is shown on the right, and the alternative of visualizing original data or, for instance, quality control measures, is shown on the left. The three studies reported in this paper, the “humanlipidome”, the “cancerlipidome”, and the “liverlipidome”, respectly, are analysed by following the routes on the left, middle and right.

Two special cases of the workflow are the visualization of associations to a categorical explanatory variable and the visualization of observations that do not come from a regression model. The first case is analyzed with the analysis of covariance (ANCOVA)^11^ and consecutive pairwise post-hoc comparisons. The ANCOVA workflow can be computed using the ‘compute_F_test_with_limma()’ and ‘compute_post_hoc_tests_with_limma()’ functions as middle-steps between Steps 4 and 5 of the main workflow (Figure 1, right; see Section 2.4 for an example). The second case is visualized by, after Step 3, directly supplying the observations to the ‘heatmap_lipidome()’ function, which creates a lipidomeR heatmap of the supplied values. Typical use cases for the ‘heatmap_lipidome()’ function are the visualization of original measurements (see Section 2.2 for an example), or the visualization of quality control measures.

Next, we test the lipidomeR with data from three publicly-available lipidomics studies. In these analyses, we give a demonstration of how to use the lipidomeR and how to interpret the results figures that the package creates. For each study, we provide a fully reproducible pipeline for obtaining these results, starting from acquiring the data and ending with final edits to the results figures.

### 2.2 The Human Lipidome

In the first experiment, we visualize the data from the 2010 hallmark study of human plasma standard reference material^12^ that, for the first time, reported the average concentrations of 535 lipids in human plasma.

Out of the lipids that were provided in the deposited data, 403 lipids were presented with an enumeration of the number of acyl-chain carbon atoms and double bonds, which is a pre-requisite for a lipidomeR heatmap. These lipids belong to 23 classes with most species in ceramides (Cers, N = 43), hexocyl-ceramides (HexCers, N=43) and sphingomyelins (SMs, N = 40).

The lipid concentrations from the human plasma standard reference are shown in Figure 2. Highest concentrations are found in cholesterol esters (CEs), phosphatidylcholines (PCs) and triacylglycerols (TGs), which emerge in red color in the respective heatmaps (see the “CE”, “PC” and “TG” panels in Figure 2). In all these three classes, mid-sized mid-unsaturated species, shown in the middle-section of the respective heatmaps, are in highest concentration. CE(18:2) is the highest-concentrated lipid, as indicated by the deepest red color in the coordinates x = 18 and y = 2 of the CE panel.

**Figure 2:**
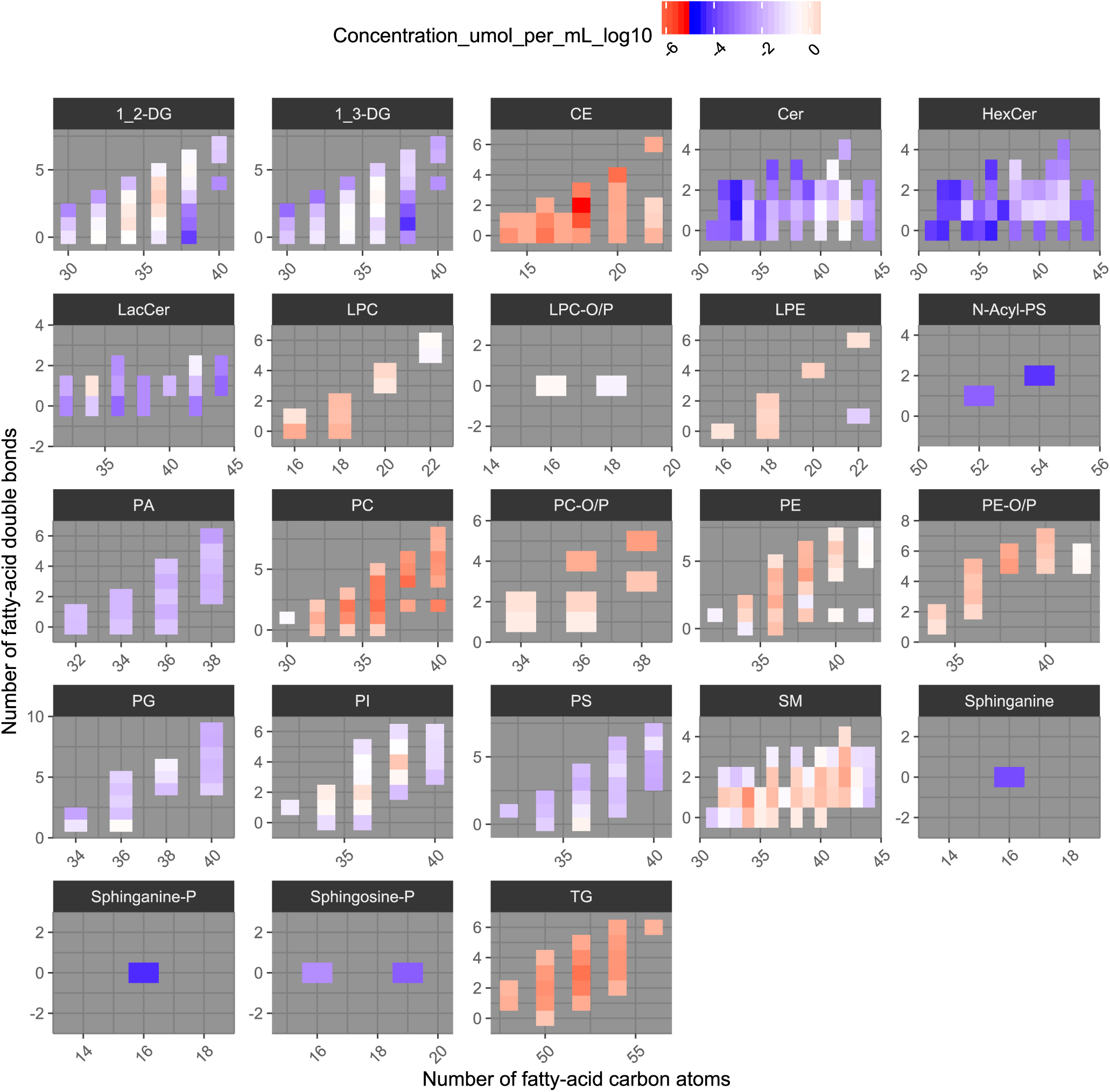
Concentrations of 403 lipids in the human plasma standard reference material. In the heatmap, each lipid is shown as a rectangle, where the color shows the respective concentration (µmol/mL in log10-scale; blue: low; white: median; red: high). The lipids are grouped into lipid classes, which are shown in distinct panels and titled according to the abbreviation of the respective lipid class. Within each lipid class, the individual lipid species are organized according to the size (x-axis) and the level of unsaturation (y-axis) of the lipid. Gray areas in the heatmap represent combinations of size and unsaturation, where no lipid was detected. For instance, the cholesterol-ester CE(18:2) is shown in the CE-titled panel at coordinates x=18 and y=2 and it is colored in deep red, since it is the highest-concentrated lipid in the data.

### 2.3 Breast Cancer Tissue

Next, we analyzed lipidomic differences between tissue from breast cancer and benign breast tumor, allowing us to test also lipidomeR’s statistical inference functionality and the integration of statistics results over the visualization of the lipidome. In this study^13^, a total of 118 tissue samples were available from 48 benign tumors, 66 malign (i.e., cancerous) tumors and 4 metastases. Out of the 773 measured lipids, 409 were unique and possible to map onto a lipidomeR heatmap in a meaningful way. These lipids belong to 13 classes. The largest lipid classes are the triacylglycerols (TGs), phosphatidylcohlines (PCs) and phosphatidylethanolamines (PEs) with 89, 67 and 54 species, respectively.

The vast majority of the lipids turn out to differ between the malign and benign tumors (Figure 3): Most consistent differences are in cholesterol-esters (CEs), lyso-phosphatidylcholines (lysoPCs) and phosphatidyl-inositols (PIs), where the lipid level is higher in cancer in all of the statistically significant differences (see panels CE, lysoPC and PI in Figure 3). Other very consistently-differing lipid classes are the diacyl-glycero-phosphoglycerols (PGs; elevated in cancer), plasmenyl-phosphatidylcholines (plasmenyl-PCs; elevated) and plasmenyl-phosphatidylethanolamines (plasmenyl-PEs; reduced). In these classes, only one of the species with a significant difference is inconsistent with the class-wide pattern.

**Figure 3:**
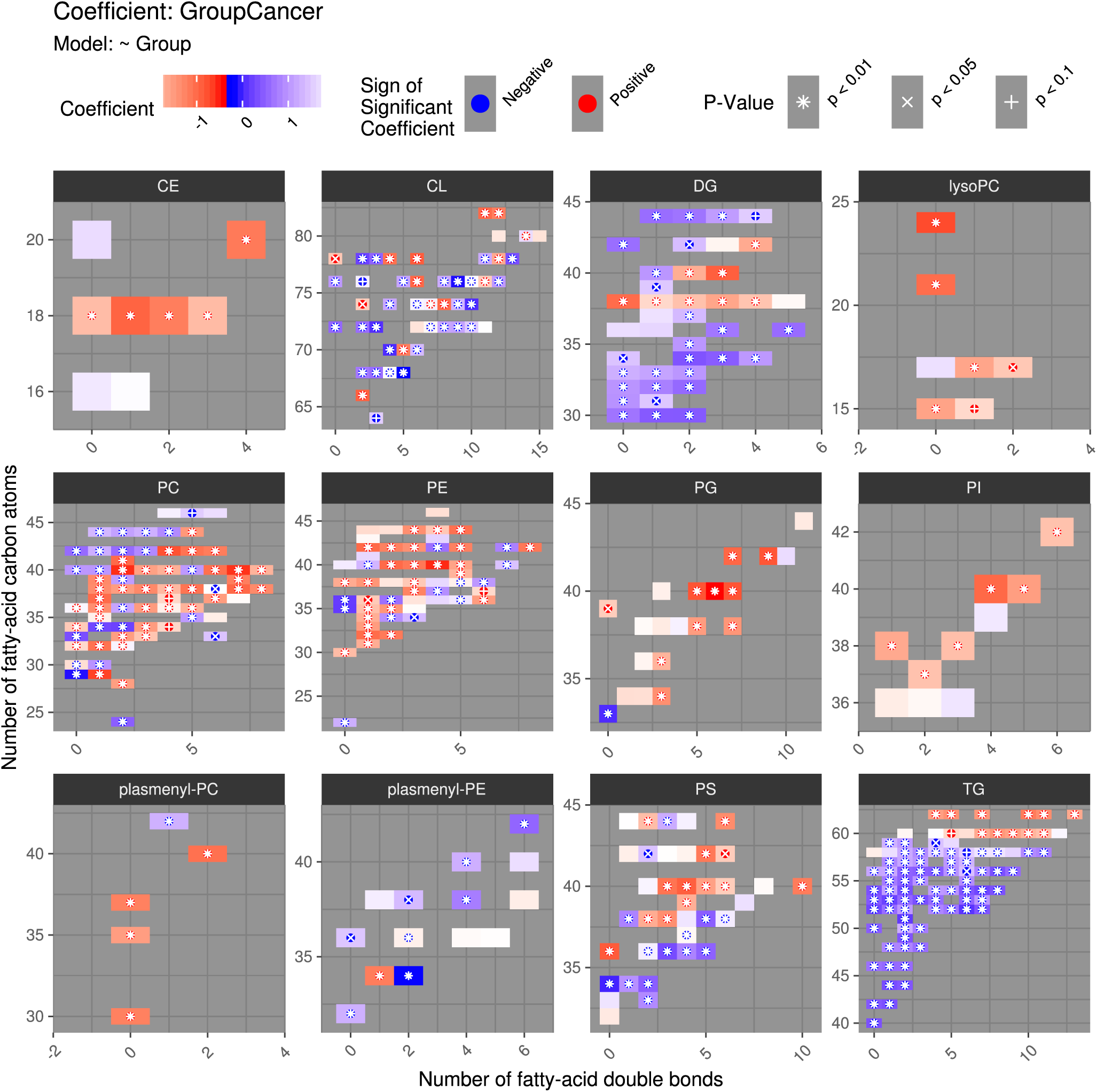
Lipidomic differences between malignant breast tumor (i.e., cancer) and benign breast tumor. In the heatmap, each lipid species is shown as a rectangle, where the color shows the difference (red: higher in cancer; white: no difference; blue: lower in cancer). The lipids are grouped by lipid class as panels, where the abbreviation of the lipid class is shown in the title of the panel. Within each class, the individual lipid species are organized according to the size (y-axis) and the level of unsaturation (x-axis) of the lipid. Lipids with statistically significant difference between cancer and benign tumor are highlighted with a character. For instance, diacyl-glycero-phosphoglycerol PG(40:6) is shown in the “PG” panel in the coordinates x = 6 and y = 40. This species is the most highly-elevated lipid among the measured lipidome in breast cancer (p < 0.01), as indicated by a deep red color and the asterisk character.

Arguably the most interesting pattern is observed in the TGs, where compounds with 59 or fewer carbon atoms in the fatty acid chains are consistently reduced in cancer while the largest compounds with 60 or more carbons were elevated (see the TG panel of Figure 3 at y ≤ 59 and y > 59, respectively). Interesting patterns are also observed in PCs, diacylglycerols (DGs) and phosphatidyl-serines (PSs): Although the majority of PCs are elevated in cancer, a cluster of large compounds are reduced (see the top-left corner of the PC panel in Figure 3). On the other hand, DGs are mainly reduced in cancer but, specifically, 38-carbon DGs break this rule with a consistent elevation (see the DG panel of Figure 3 at y = 38). This elevation also extends to few poly-unsaturated DGs with 40 or 42 carbon atoms (see the DG panel at y = 40 and 42). In phosphatidyl-serines (PSs), a cluster of mid-sized poly-unsaturated compounds are elevated while the smaller compounds are reduced (see the PS panel of Figure 3). The remaining lipid classes, namely, cardiolipins (CLs) and phosphatidyl-ethanolamines (PEs) are mixed with elevated and reduced lipids with no apparent pattern related to size or saturation.

In this demonstration, we reported differences between cancerous and benign tumor samples. The small group of metastatis samples differed with similar but somewhat stronger patterns as the cancer tumor samples, when compared to the benign tumor samples. These additional results are shown in Supplementary Material, p.5.

### 2.4 The Spectrum of Nonalcoholic Fatty Liver Disease

Finally, we investigate a large study^14^ on non-alcoholic fatty liver disease (NAFLD), allowing us to demonstrate more complex statistical analysis and interpretation of the results. The data from the study include lipidomics of liver, plasma and urine samples from 88 participants with NAFLD ranging between healthy (N=31), steatosis (N=17), hepatosis (N=20) and cirrhosis (N=20). Out of the 542 measured lipids in the liver, 383 are unique and possible to map onto a lipidomeR heatmap in a meaningful way. These lipids belong to 31 classes. The largest lipid classes are the triacylglycerols (TGs), 1,2-diacylglycerols (DGs) and free fatty acids (FAs) with 90, 4 and 25 species, respectively.

In liver tissue, 144 lipids have a difference between the NAFLD categories in the analysis of covariance (ANCOVA; p < 0.05; Supplementary Material, p.8). In post-hoc comparisons, which we next describe as the main result of this experiment, it becomes clear that steatosis and hepatosis exhibit largely similar lipidomic patterns while cirrhosis is markedly different (Figure 4).

**Figure 4:**
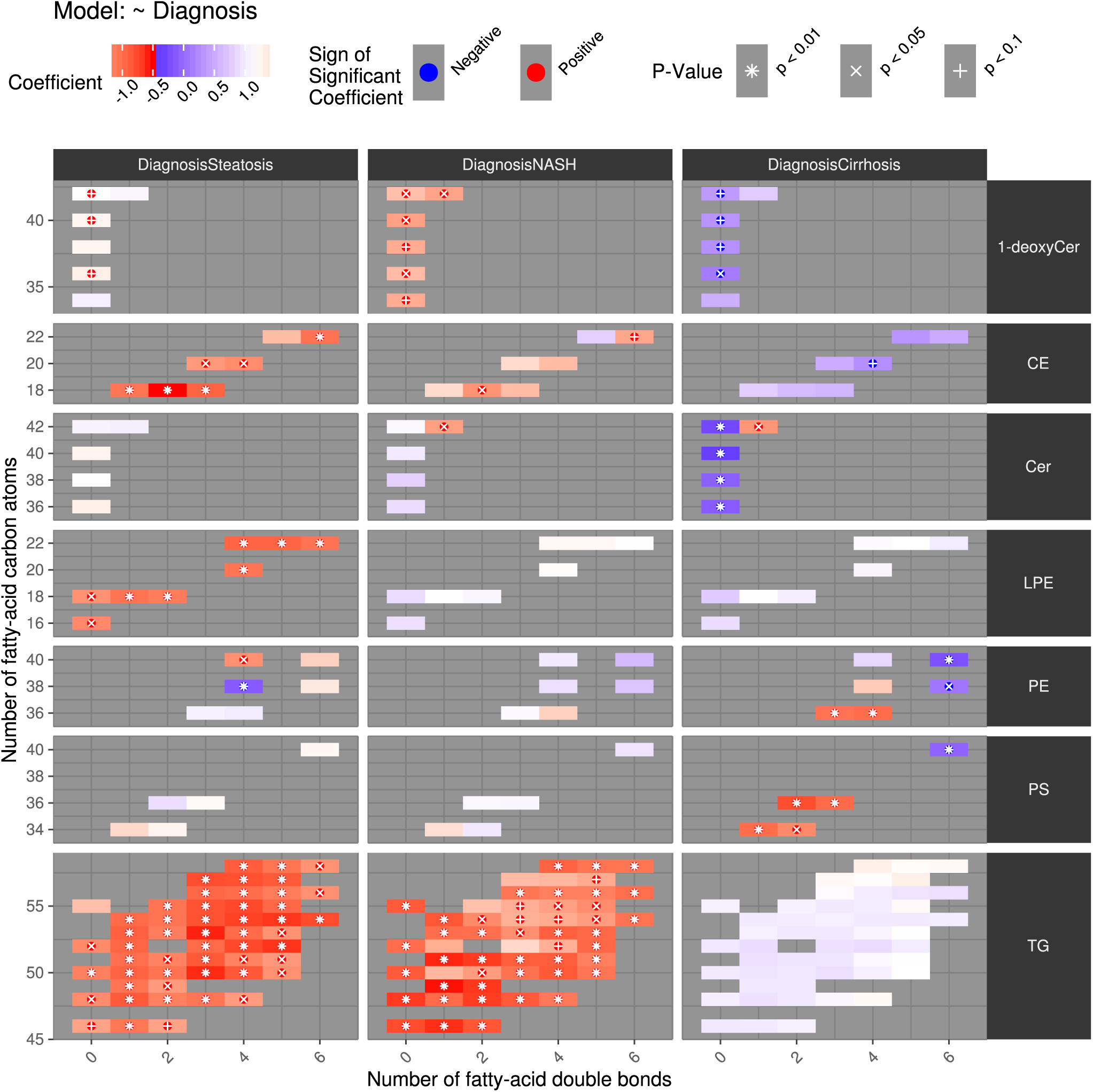
Lipidomic differences between three stages of non-alcoholic fatty liver disease (NAFLD) and healthy control. In the heatmap, each lipid species is shown as a rectangle, where the color shows the difference (red: lower in healthy control; white: no difference; blue: higher in healthy control). The lipids are grouped by disease stage (panel columns) and lipid class (panel rows). The lipid class abbreviation is shown in the row panel title (right) and the disease stage is named in the column panel title (top). Within each panel, representing a pair of lipid class and disease stage, the individual lipid species are organized according to the size (y-axis) and the level of unsaturation (x-axis) of the lipid. Lipids with statistically significant difference are highlighted with a character. For instance, the cholesterol-ester CE(18:12) is shown in the row “CE” in the coordinates x = 2 and y = 18. The lipid is highly elevated in steatosis (p < 0.01), as indicated by a deep red color and the asterisk character in the “DiagnosisSteatosis” column. The same lipid is still elevated in NASH (p < 0.05), as indicated by a red color and the cross character in the “DiagnosisNASH” column. Finally, CE(18:2) is no longer elevated in cirrhosis but rather is slightly in lower level than in the control group (not significant), as indicated by a mild blue color in the “DiagnosisCirrhosis” column.

Steatosis – the first stage of NAFLD – is defined by an accumulation of triacylglycerols (TGs) in the liver. This is also observed in the data as virtually every measured unsaturated TG species is elevated (see the bottom-left panel the “DiagnosisSteatosis” column and in the TG row of Figure 4, where rectangles at location x > 0 appear red with an asterisk or cross, indicating statistical significance). In most of the saturated TGs, though the elevation is not significant (see the rectangles at location x = 0). The most marked increase appears in mid-sized unsaturated TGs (see the rectangles at location y = 50 and x = 3 as well as at y = 53 and y = 3, in corresponding to TGs 50:3 and 53:3, appear in most intense red, indicating a strong elevation in steatosis).

Beyond the TGs, steatosis is characterized by elevated 1,2-diacylglycerols (1,2-DGs), cholesterol-esters (CEs) and lyso-phosphatidyl-ethanolamines (LPEs), as indicated by the red color in the panels at the “1,2-DG”, “CE” and “LPE” rows of at the “DiagnosisSteatosis” column. In CEs and LPEs, the elevation appears throughout the board, albeit in LPEs the elevation appears as somewhat stronger in non-saturated lipids (see rectangles in the LPE-Steatosis panel with x > 0. In 1,2-DGs, the elevations are weaker with only two 34-carbon 1,2-DGs at p < 0.05 (see the rectangles in the 1,2-DG panel with y = 34). In other lipid classes, only minor or inconsistent changes are observed in steatosis.

The second disease stage – the non-alcoholic steatohepatitis (NASH) – is also defined by an accumulation of TGs in the liver. However, the elevation profile is different from steatosis: The most elevated TGs in NASH are the saturated TG(57:0), and the smaller TGs 46:1, 49:1 and 51:1 (see the NASH-TG panel of Figure 4 at locations x = 57 and y = 0 as well as x = 1 and y = 46, 49 and 51). In contrast, many large or mid-sized and unsaturated TGs are only mildly elevated (p > 0.05). Beyond the TGs, the same pattern of elevated CEs and LPEs is not seen as in steatosis.

The terminal stage of liver disease – the cirrhosis – is completely different from NASH and steatosis. Remarkably, there is no difference in the profile of TGs between a cirrhotic and a healthy liver (see the panel in the DiagnosisCirrhosis column and in the TG row). Instead, a cirrhotic liver is characterized by a decrease in saturated ceramides (Cers; p < 0.01; see the DiagnosisCirrhosis-Cer panel at x = 0). Indicative but considerably weaker decrease is observed in saturated 1,2-DGs (see the DiagnosisCirrhosis-1,2-DG panel of Figure 4). Finally, the most unsaturated PEs and phosphatidyl-serines (PSs) are reduced in cirrhosis (see the panels at the PE and PS rows and at the DiagnosisCirrhosis column at x = 6 in the middle-right of Figure 4). In smaller and more saturated PEs and PSs, the aberration is opposite with an elevation in cirrhosis (see the same panels at x = 1…3).

Beyond these consistent changes during the progression of NAFLD, changes in other lipid classes, such as sphingomyelins (SMs), are observed (Supplementary Material, p.10). While these patterns also vary across the disease stages, all the lipid classes that were not presented in Figure 4 had fewer than four changes in all the stages combined, indicating a lower consistency of the aberrations than observed in 1,2-DGs, CEs, Cers, LPEs, PEs, PSs and TGs.

## 3 Discussion

The NIDKK/NIST reference data consisted of only one concentration value per lipid. Thus, inter-individual variation cannot be assessed based on these data. Despite this limitation, the data provide insight into, which lipids classes are abundant in human plasma, and how much the levels of individual lipid species within a class differ. In this paper, we provided a systematic visualization of the human lipidome. We argue that this visualization created additional value to the published data by uncovering detailed lipid concentration patterns in human plasma.

Investigation of breast tumors revealed a lipidome-wide dysregulation in the cells’ lipid balance in cancer. The lipidomeR heatmap visualization helped uncover a size-and-saturation-dependent pattern in DGs, PCs and, particularly, in TGs: The largest TGs were elevated in cancer although the bulk of TGs were clearly reduced.

Analysis of the non-alcoholic fatty liver (NAFLD) study revealed distinct lipidomic patterns of the disease stages. While the major differences were reported in the original publication of the study^14^, we argue that the analysis presented here delivered a significantly new interpretation of the data and presented the data in a systematic manner that allows the audience to understand the big picture in these complex data. Particularly, the ANCOVA-post-hoc procedure coupled with interpretable heatmaps allowed for the description of disease-stage-specific patterns, which could have clinical application.

We believe that the lipidomeR can improve the interpretability and reproducibility of lipidomics research. It is our aim to demonstrate more use cases of this tool in new translational studies in medical research, as already has been done in Tofte, et al., 2019^15^ and Al-Sari, et al., 2020^16^. We aim to make additional functionalities available to the lipidomeR by extending the portfolio of suppoted data analyses to a broad set of analysis cases in clinical and biomedical research. Some of these planned additions have already been developed, including the integrated computation and visualization of other regression models, such as mixed-effect models (see Ahonen, et al., 2019^17^) and time-to-event models (see Tofte, et al., 2019), and novel integrative visualizations, such as bipartite graphs (see Tofte, et al., 2019b^18^) and partial correlation networks (see Trost, et al., 2020^19^). Finally, by providing reproducible examples directly in the package, we aim to making the tool accessible to a broad set of users with diverse academic backgrounds, which is a typical feature of metabolomics research.

## 4 Methods

Publicly available datasets from the Metabolomics Workbench were analysed and visualized to demonstrate the method.

Data were downloaded from the Metabolomics Workbench in the mzTab format. Consecutive work was done in R. First, a dataset-specific function to load and prepare the data was written to extract observations and experiment design from the mzTab data.

### 4.1 The lipidomeR Package

Once loaded into a compatible format, the data were analysed and visualized with the R-package lipidomeR, which can be installed in R (R>3.5.0) with the command ‘install.packages(“lipidomeR”)’. The package can also be found at https://cran.r-project.org/web/packages/lipidomeR/.

The steps from ready data to result are as follows: (1) Define the list of lipid names in the data. (2) Parse the lipid names into mappable values describing the lipid class, the number of carbon atoms in the fatty acid chain (i.e., the size of the lipid) and the number of double-bonds in the fatty acid chain (i.e., the degree of unsaturation). This step is done by calling the ‘map_lipid_names()’ function. (3) If applicable, compute lipid-wise regression models to describe the data in terms of the experimental covariates. This step is done by calling the ‘compute_linear_models_with_limma()’ function. (4) Finally, visualize the fitted regression model coefficients on a heatmap that shows the measured lipidome, grouped by the lipid class and experimental covariate, and organized according to the size and the level of unsaturation of the lipids. This step is done by calling the ‘heatmap_lipidome_from_limma()’ function.

### 4.2 The Human Lipidome

The data for this study were downloaded from the project ST000004 at the Metabolomics Workbench (http://dx.doi.org/10.21228/M8MW26). The script for preparing the data is available at https://github.com/tommi-s/lipidomeR/blob/master/scripts/prepare_humanlipidome.md.

The data set contains only one concentration value per lipid. That is, there are no replicate measurements reported but the values are from a single measurement of the reference material, which itself is a pooled composite of plasma from an order of hundred healthy individuals. Due to the N=1 nature of the study, statistical models were not fitted to these data, meaning that the Step 3 in the lipidomeR workflow (described in Section 4.1) was not applied. Instead, the lipid concentration values from the data were directly used in the lipidomeR heatmap visualization at Step 4.

The prepared data set is available in the lipidomeR package under the name ‘humanlipidome’. The results reported in this paper can be reproduced by running the example in the documentation of the ‘humanlipidome’ data set. The example can also be viewed on browser at https://lipidomer.org.

### 4.3 Breast Cancer Tissue

The data for this study were downloaded from the project ST001111 at the Metabolomics Workbench (http://dx.doi.org/10.21228/M8RX01). The script for preparing the data is available at https://github.com/tommi-s/lipidomeR/blob/master/scripts/prepare_cancerlipidome.md.

In the interest of clarity for this experiment, we focused on the comparison between malignant tumor (cancer) samples and benign tumor samples. The association between the tumor diagnosis and lipid levels was analysed with lipid-specific linear regression models, where the diagnosis category entered the model as categorical independent variable and one lipid feature at a time as a dependent variable.

The inferred model coefficients were visualized in a lipidomeR heatmap. Particularly, a heatmap to visualize the difference between the malignant and benign tumor samples was created. To improve readability of the figure by making the number of sub-panels fit a 3-by-3 grid, the lipid class of glycerophosphates (PAs) was omitted from the figure. Among the lipid classes, PAs were chosen on the ground that the class is rather highly scattered in terms of the individual species size and saturation, indicating a low coverage of the entire PA class. Figures including also the PAs were also produced for Supplementary Material.

The prepared data set is available in the lipidomeR package under the name ‘cancerlipidome’. The results reported in this paper can be reproduced by running the example in the documentation of the ‘cancerlipidome’ data set. The example can also be viewed on browser at https://lipidomer.org.

### 4.4 The Spectrum of Nonalcoholic Fatty Liver Disease

The data for this study were downloaded from the study ST000915 of the project PR000633 at the Metabolomics Workbench (http://dx.doi.org/10.21228/M8V961). In the interest of clarity of demonstration for this experiment, we focused on lipidomic data from the liver samples. The script for preparing the data is available at https://github.com/tommi-s/lipidomeR/blob/master/scripts/prepare_liverlipidome.md.

Each lipid species was tested for any difference between the NAFLD categories with analysis of covariance (ANCOVA) by using lipidomeR. The lipids passing the ANCOVA with a significant difference were further investigated with post-hoc tests comparing the healthy group to the other NAFLD groups. The post-hoc comparisons between the disease stages of steatosis, non-alcoholic steatohepatitis (NASH) and cirrhosis against healthy control were visualized as lipidomeR heatmaps. Lipid classes with five or more aberrations with statistical significance (p < 0.05) were shown in the heatmap in Figure 4 while a heatmap with all lipids were also produced for supplementary material (Supplementary Figure N).

The prepared data set is available in the lipidomeR package under the name ‘liverlipidome’. The results reported in this paper can be reproduced by running the example in the documentation of the ‘liverlipidome’ data set. The example can also be viewed on browser at https://lipidomer.org.

## Data Availability

All data presented in this paper are available as part of the lipidomeR package. Also the code to reproduce the results are available as examples in the lipidomeR package as well as in the Supplementary Material and at https://lipidomer.org. The original data sets are available at the Metabolomics Workbench repository. Scripts for preparing the data from the original data sets are available at https://github.com/tommi-s/lipidomeR/tree/master/scripts. See the Methods Section for details.

## Acknowledgement

We thank the authors of the human lipidome study^12^, the authors and donors of the breast cancer study^13^ and the authors and donors of the NAFLD study^14^ for providing their data with an open access. We thank the Metabolomics Workbench^20^ for providing a platform for open-access metabolomics data, where also these lipidomics data sets had been deposited. We thank the R-project for providing the infrastructure for hosting the lipidomeR package.

## Author Contributions

T.S. designed and implemented the tool, planned and carried out the experiments, interpreted the results, and wrote the manuscript. All authors critically reviewed the manuscript.

